# Tree growth enhancement drives a persistent biomass gain in unmanaged temperate forests

**DOI:** 10.1101/2022.11.16.516717

**Authors:** Laura Marqués, Ensheng Weng, Harald Bugmann, David I. Forrester, Brigitte Rohner, Martina L. Hobi, Volodymyr Trotsiuk, Benjamin D. Stocker

## Abstract

While enhanced tree growth over the last decades has been reported in forests across the globe, it remains unclear whether it drives persistent biomass increases of the stands, particularly in mature forests. Enhanced tree growth and stand-level biomass are often linked with a simultaneous increase in density-driven mortality and a reduction in tree longevity. Identifying empirical evidence regarding the balance between these processes is challenging due to the confounding effects of stand history, management, and environmental changes. Here, we investigate the link between growth and biomass via the shift in the negative relationship between average tree size and stand density (tree number). We find increasing stand density for a given tree size in unmanaged closed-canopy forests in Switzerland over the past six decades and a positive relationship between growth and stand density - qualitatively consistent with simulations by a mechanistic, cohort-resolving ecosystem model (LM3-PPA). Model simulations show that, in the absence of other disturbances, enhanced growth persistently increases biomass stocks despite simultaneous decreases in carbon residence time and tree longevity, independent of assumptions about the drivers of tree mortality. However, the magnitude of simulated changes critically depends on the shape of the mortality parameterizations. Our analyses reconcile reports of growth-induced reductions of tree longevity with model predictions of persistent biomass increases, and with our finding of a trend towards denser forests in response to growth - also in mature stands.

## 1. Introduction

Vegetation demography processes, namely growth, recruitment, and mortality are being altered by global environmental change (McDowell *et al*. 2020). Enhanced tree growth over the last decades has been widely reported (Cole *et al*. 2009; McMahon *et al*. 2010; Fang *et al*. 2014; Wu *et al*. 2014; Brienen *et al*. 2015). Trends in growth and forest functioning have been attributed to increased nutrient inputs by atmospheric deposition, rising temperatures and extended growing seasons (Pretzsch *et al*. 2014; Anderegg *et al*. 2015), and elevated atmospheric carbon dioxide (eCO_2_) (Huang *et al*. 2007; Lewis *et al*. 2009; Phillips *et al*. 2009; Pan *et al*. 2011; Hubau *et al*. 2020). Also, biomass stocks have been reported to have increased in forests around the globe (Pan *et al*. 2011), unless large-scale disturbances and extreme events reversed long-term trends (Wang *et al*. 2021). However, it remains debated to what extent increased biomass stocks are a consequence of accelerated tree growth in response to environmental change or of recovery from past disturbances and land use (Frelich 2002; Gloor *et al*. 2009). Disturbance history and stand age are dominant factors determining forest biomass stocks (Bradford *et al*. 2008) and can mask the effects of environmental change. This limits our understanding and poses an observational challenge for attributing the observed forest carbon (C) sink (Pan *et al*. 2011) to anthropogenic versus environmental drivers, and for answering the question of whether accelerated tree growth, induced by environmental change, leads to persistent increases in forest biomass stocks.

Direct evidence for environmental change effects on growth and biomass comes from ecosystem manipulation experiments. Free Air CO_2_ Enrichment (FACE) experiments have identified increases in biomass production in response to enhanced CO_2_ (Ainsworth & Long 2005; Norby *et al*. 2005; Hovenden *et al*. 2019; Jiang *et al*. 2020; Walker *et al*. 2021). However, positive effects on biomass have been argued to be transitory (Bugmann & Bigler 2011; Büntgen *et al*. 2019; Fatichi *et al*. 2019; Fleischer *et al*. 2019; Brienen *et al*. 2020), limited to young forests (Norby & Zak 2011), and absent in mature forests (Jiang *et al*. 2020). This argument can be linked to two hypothesised mechanisms. First, the *progressive nitrogen limitation* hypothesis (Luo *et al*. 2004) states that soil N gets progressively depleted as biomass stocks accumulate. By implication, old-growth forests are prone to N scarcity, reducing growth and triggering additional feedback via increases in the C:N ratio of litter and ensuing decreases in net mineralisation rates. Second, the *grow-fast-die-young* hypothesis (hereafter GFDY) (Bugmann & Bigler 2011; Körner 2017; Büntgen *et al*. 2019; Brienen *et al*. 2020) posits a reduced longevity of fast-growing trees, as described in more detail below. Both hypotheses predict that a positive response of biomass stocks to enhanced growth would be reduced or fully suppressed in mature stands. Indeed, the first FACE experiment conducted in a mature stand did not show increased carbon sequestration at the ecosystem level (Jiang *et al*. 2020), even if it is not fully established whether the observed response was due to forest demography or nutrient availability-related mechanisms (Ellsworth *et al*. 2017).

The GFDY may result from the evolution of species’ life-history strategies along the resource conservation vs. exploitation spectrum, leading to fast-growing and short-lived species at one end (mostly pioneers in forest succession), and slow-growing and long-lived species at the other end. The GFDY trade-off could also be the outcome of forest demography processes leading to a reduction of carbon residence time when tree growth is enhanced over time across individuals in a stand. While much empirical support for the GFDY hypothesis is based on variations across species (Loehle 1988; Wright *et al*. 2010; Brienen *et al*. 2015), the emergent negative feedback between growth and biomass changes has also been argued to govern the response of forest stands to environmental change in the absence of effects by species replacement (Bugmann & Bigler 2011; Brienen *et al*. 2020). Growth-longevity trade-offs within species have been found previously (Bigler & Veblen 2009; Büntgen *et al*. 2019). The mechanisms underlying the negative feedback at the forest stand scale relate to competition for light and the tree’s C balance. Accelerated growth and crown expansion under a constant canopy space constraint (Zeide 1993) can drive the exclusion of short trees from the canopy, intensifying competition for light, and potentially enhancing their mortality. Consequently, accelerated growth can speed up the tree’s life cycle through earlier mutual shading in a closed forest stand. Faster growth can also lead to earlier attainment of a critical tree size where hydraulic, mechanical, or C balance constraints (such as insufficient investment into defence) pose limits to further growth and may trigger mortality (Collalti *et al*. 2020; McDowell *et al*. 2022).

Allometric relationships of tree diameter, height, and crown area, combined with the packing constraint, lead to an emergent relationship between average tree size or biomass and the number of trees per unit area in closed-canopy forests. For monospecific and even-aged stands, this relationship has been described by a power-law relationship of the number and quadratic mean diameter of trees in a closed forest stand - Yoda’s Law (Yoda 1963). Forest data following Yoda’s Law align along the so-called self-thinning line (STL) - the linear form of the tree number vs. mean tree size relationship in a double-logarithmic plot. As a forest stand matures, the increase in tree size and the simultaneous decrease of tree number, i.e., density-driven mortality, is determined by the intercept and slope of the STL. The use of STLs has a long tradition in forest management (West *et al*. 1997; Enquist *et al*. 1998) and research (Pretzsch 2006; Charru *et al*. 2012). Prescribed, site-specific and temporally stationary STLs have been used in forest demography models (Mäkelä *et al*. 2000) for simulating forest stand dynamics and density-driven mortality, subject to the packing constraint (Landsberg & Waring 1997).

By implication of a stationary STL, accelerated growth of trees will simply lead to their accelerated progression along the constant STL - consistent with the GFDY hypothesis. Hence, the relative change in biomass stocks should be negligible, irrespective of the relative enhancement of growth, because total stand-level biomass is largely constant along the STL. However, the position of the STL is a reflection of site quality (climate and soil properties) and is affected by species identity (Forrester *et al*. 2021). Although temporal changes in the STL are still in doubt (Pretzsch *et al*. 2014), a recent study found that the STL shifted such that forest stands were able to carry higher biomass stocks in a CO2 fertilization experiment (Kubiske *et al*. 2019). Thus, analyzing the changes in the self-thinning trajectories is a sound way to tackle the GFDY hypothesis. To follow this approach, data from forest inventories are particularly valuable, even though many forest inventory plots are affected by prior management and cannot be assumed to have reached maturity and, consequently, steady-state biomass stocks. Still, long-term observations from unmanaged closed-canopy forest plots offer a unique opportunity to investigate growth-longevity trade-offs driven by self-thinning and environmental changes.

Global Dynamic Vegetation Models (DGVMs) are widely used for simulating the response of the terrestrial C cycle to global environmental change (Arora *et al*. 2019; Friedlingstein *et al*. 2021). However, these models have traditionally relied on simplifications of forest stand dynamics by resolving only an average individual tree (Sitch *et al*. 2003; Fisher *et al*. 2019), thus not mechanistically accounting for size-dependent light competition and mortality (Purves *et al*. 2008; McDowell *et al*. 2011; Evans 2012; Bugmann *et al*. 2019). Therefore, such models are not suitable to investigate mechanisms underlying the GFDY hypothesis. This simplification may also imply unrealistic simulations of growth-biomass relationships under environmental change (Friend *et al*. 2014; Yu *et al*. 2019; Pugh *et al*. 2020). For example, a constant background mortality specified in models (Bugmann *et al*. 2019) may imply an overestimation of the relative change in biomass stocks per unit relative change in growth (*constant turnover rate*, Box 1), or a constant prescribed STL (Landsberg & Waring 1997) may imply an underestimation of the same (*constant self-thinning*, Box 1).

### Box 1.

#### Approach to link changes in growth, biomass and stand density

Mechanistic models (e.g., DGVMs) represent the carbon cycle in terrestrial ecosystems as a cascade of pools and fluxes (Randerson *et al*. 1997; Smith *et al*. 2013). The assumption that pool-specific residence times are independent of input fluxes and pool sizes, combined with constant relative allocation of fluxes to downstream pools and respiration, leads to linear systems dynamics (Luo & Weng 2011; Xia *et al*. 2013) and a proportional scaling of fluxes and pools with the ultimate C input flux to the system - CO_2_ assimilation by photosynthesis. Density-driven mortality is a process that introduces a negative relationship between biomass C residence times and biomass production, thus triggering a feedback which leads to a deviation from linear systems dynamics. The analysis presented here is designed to diagnose this deviation from model outputs by investigating relative changes in biomass (*B*) and growth (*G*).

Let us consider the wood C pool size, i.e., biomass (*B*) corresponding to the difference between growth (*G*) and mortality (*M*):

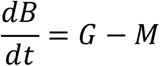

The wood C pool dynamics can also be described by first-order kinetics, such as:

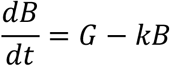

With 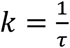 describing the carbon turnover rate, i.e., the inverse of carbon residence time. At the steady-state, 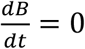, and thus the wood C pool size is equal to:

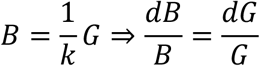

Three cases can be distinguished and have different implications for changes in the self-thinning relationship in response to changes in *G*.

1. *Constant turnover rate*, 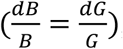, which implies that a relative enhancement of G leads to an equal relative enhancement of B and a shift upwards in the STL. This linear response can also be seen when carbon residence time is modelled as a function of the climate or prescribed disturbances even if effective residence time change.
2. *Accelerated turnover*, 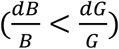, an intermediate case which leads to a non-linear response where the relative increase in B is smaller than the relative increase in G. Carbon turnover time is reduced but an upward shift in the STL is still observed.
3. *Constant self-thinning*, (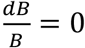 in case of constant B along the STL), which implies that G enhancement accelerates tree life cycle to a degree that nullifies the change in B, as underlined in the GFDY hypothesis. Trees progress faster along the STL, but the position remains unchanged.

Considering this link between biomass and the STL, cases *i* and *ii* are marked with an upward shift of the STL, whereby the upward shift is larger for *i* than for *ii*. Investigating STL changes over time and in relation to variations in *G* thus yields insight into the (essentially unobservable but key) steady-state *G*-*B* relationship.

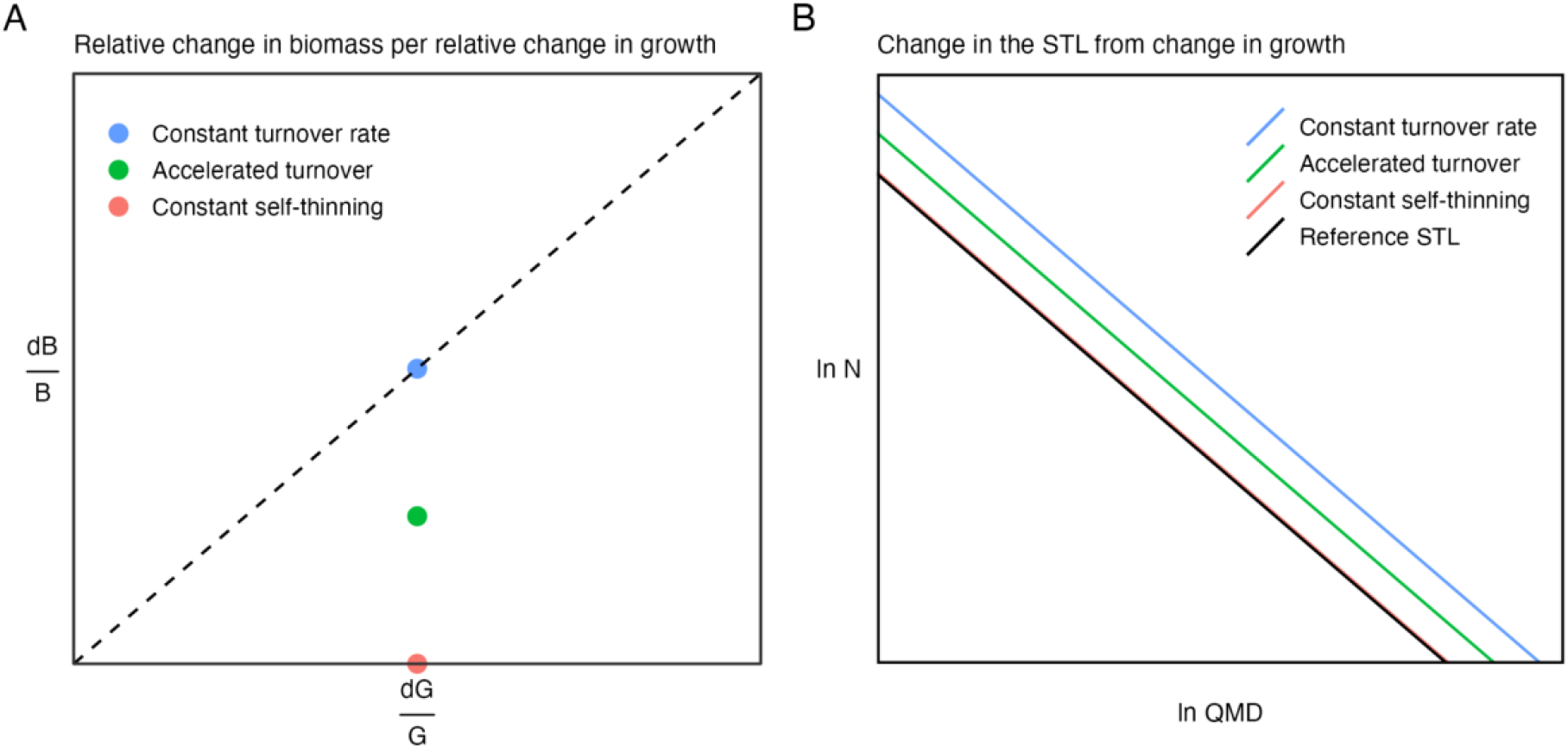
Conceptual model of biomass and STL responses to growth enhancement. (A) Responses of biomass stocks where an enhancement in growth may lead to no biomass increment (*constant self-thinning*, red circle) or to equal relative biomass increment (*constant turnover rate*, blue circle in the dashed 1:1 line). (B) Responses of the STL where growth enhancement may lead to no changes in the STL (*constant self-thinning*, red line on top of the lower solid black line, considered as a reference STL) or to a shift upwards in the STL (*constant turnover rate*, blue line). The intermediate stages of these two extreme assumptions representing an *accelerated turnover* are shown by a green circle (A) and line (B).

Here, we evaluated empirical and theoretical support for the GFDY hypothesis by investigating observed and simulated changes in the relationship between growth and biomass, and the underlying mechanisms in mature forest stands. We used long-term forest data from unmanaged stands in Switzerland to evaluate whether the STL has shifted through time and whether variations in the STL are influenced by stand-level growth. Then, using a vegetation demography model (LM3-PPA, Weng *et al*. 2015, 2019) that combines a treatment of tree-level physiology and C balance with the simulation of competition for light and mortality, we explored the underlying mechanisms and investigated under which conditions and model formulations persistent biomass stock increases result from growth enhancements.

## 2. Material and methods

### 2.1. Observed forest trends

#### 2.1.1. Forest data

Inventory data from mixed forests were obtained by combining observations from three sources: the Swiss National Forest Inventory (NFI) (Fischer & Traub 2019), the Experimental Forest Management (EFM) network (Forrester *et al*. 2021), and the Swiss Natural Forest Reserves (NFR) (Hobi *et al*. 2020). These data cover a large environmental gradient and a variety of site conditions, making Swiss forests an interesting study case. From the compiled dataset, we selected unmanaged plots based on relevant information specific to each original dataset. We selected NFI plots free of management for at least 70 years. This information was based on standardized interviews with the forest services (Portier *et al*. 2021). We considered the EFM plots with no intervention since monitoring started, with an average of 40 years. None of the NFR plots has experienced any management since the establishment of the forest reserves, with an average of 85 years unmanaged. The combined dataset covers the period from 1933 to 2019 and features 516 plots from the NFI, 18 plots from the EFM, and 269 plots from the NFR. The measurement intervals varied between 10 and 12 years, depending on sampling design, growth rates and environmental conditions (see table S1 for more details on data characteristics).

#### 2.1.2. Stand measurements

In most cases, tree diameter at 1.3 m height (DBH, cm) was measured on all trees with DBH ≥ 4 cm (NFR), 8 cm (EFM), or 12 cm (NFI). For each stand, quadratic mean diameter (QMD, cm), stand density (N, trees ha^−1^), and total biomass (B, kg m^−2^) were calculated for each measurement year. Biomass was estimated for the EFM and NFR plots following the allometric equations described in Forrester *et al*. 2017, where biomass is predicted from DBH and stand basal area. Species-specific equations included wood density (g cm^−3^), or specific leaf area (SLA, m^2^ kg^−1^) also obtained from Forrester *et al*. 2017. The NFI dataset provided biomass estimates following the methodology described in Fischer & Traub 2019. For these plots, biomass is calculated from the estimated volume of the living trees based on tree-species-specific wood densities. Net biomass change (ΔB) was estimated as kg m^−2^ yr^-1^ by dividing the biomass difference from successive pairs of measurements by the length of the observation period. We estimated the dominant species in each stand as the one with the highest basal area (m^2^ ha^-1^). To evaluate the changes in species composition over time, we calculated the Bray-Curtis dissimilarity index (Bray & Curtis 1957) by stem number for each forest stand, which ranged from 0.14 to 0.26 (Table S1).

#### 2.1.3. Data analysis

To estimate the self-thinning relationships, a subset of plots was selected that feature high stem numbers for a given QMD. The range of QMD in the plots was divided into 30 bins of ca. 3 cm. Across all bins, we performed a sensitivity analysis to select the best cut-off criterion between the 55^th^, 75^th^, and 90^th^ percentile of plots in terms of their density within each QMD bin. The STLs were finally estimated using the 75^th^ percentile cut-off criterion, resulting in 318 plots, with measurements spanning from 1946 to 2019. This selection criterion provided a sufficient sample of plots to allow for statistically estimating changes in the size-density relationship.

The self-thinning relationships were determined by regressing the logarithms of tree density and QMD. We examined whether the STLs exhibited any shifts over time or any relationship with stand growth rate. We used Linear Mixed Models (LMMs) to evaluate how the STL depends on (i) calendar year and (ii) growth anomalies. To estimate growth anomalies, we first fitted the stand-level ΔB against QMD using a Generalized Additive Mixed Model (GAMM) to remove the size effect on biomass accumulation and extracted the residual values (Fig. S1). The general structure of the LMMs can be summarised as:

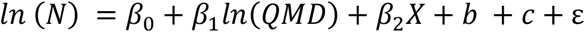

where stand density (*N*) is the dependent variable, and the fixed factors are *QMD* and *X*, which represent either calendar year or growth anomalies. The parameters *b* and *c* are the random intercepts with plot identity and dominant species as grouping variables, respectively, and *ε* is the residual error term. We did not include an interaction effect between QMD and the predictors because we were interested in the upward shift in the STL, i.e., the change in the intercept. Fixed effects selection was based on Akaike Information Criterion (AIC), which selected the models with main effects (no interactions) as the most parsimonious models (Burnham & Anderson 2003). Parameter estimation was made using restricted maximum likelihood (REML), which minimises the likelihood of the residuals from the fixed-effect portions of the model (Zuur 2009). The percentages of variance explained by the fixed and random effects of the best model were obtained according to Nakagawa & Schielzeth 2013.

All statistical analyses were performed using the R statistical software version 4.0.5 (R Core Team, 2021). We fitted the GAMM using the *gamm4* R package (Wood 2017) and the LMMs using the *lme4* R package (Bates *et al*. 2015) and calculated p-values with the *lmerTest* R package (Kuznetsova *et al*. 2017).

### 2.2. Modelling approach

#### 2.2.1. Model description

LM3-PPA is a cohort-based vegetation demography model combining leaf-level ecophysiology, individual-level competition for light and soil resources, forest structural dynamics, and biogeochemical processes (Weng *et al*. 2015). The model links a standard photosynthesis model (Farquhar *et al*. 1980; Leuning *et al*. 1995) with tree growth and allometry, and scales from the geometry of individual trees to canopy structure and competition for light using the Perfect Plasticity Approximation (PPA) (Purves *et al*. 2008). The PPA assumes that individual tree stems and crowns are organised to fill the canopy irrespective of a tree’s lateral positioning and thus form discrete canopy layers, within which all plants receive the same incoming radiation. Exclusion from the canopy and shading is determined based on a tree’s height in relation to the critical height of the canopy (*H*\*), which is defined as the height of the shortest canopy tree, whereby the crown areas of canopy trees sum up to unity (minus a constant gap fraction). LM3-PPA allows for an explicit representation of cohorts of equally sized individuals and for a treatment of a tree’s C balance and mortality. The model thus simulates size-structured competition for light, demographic processes, and dynamics of a forest stand. It has been comprehensively documented and evaluated against data from Eastern US temperate forests (Weng *et al*. 2015) and temperate to boreal forests in North America (Weng *et al*. 2017). For the present study, we used the model version described in Weng *et al*. 2019 but disabled the nitrogen limitation constraints. We re-calibrated the model for simulations representing conditions in Central European forests (see section 2.2.3).

#### 2.2.2. Mortality formulations and parameterization

To test the GFDY hypothesis, two alternative assumptions about the structural dependence of mortality (m) were defined for canopy trees (tree height above *H*\*).

A size-dependent mortality was specified for the upper canopy layer, assuming the yearly mortality rate of the upper-canopy trees to follow a power law relationship with tree’s diameter (Eq 1). In this formulation, *d* is the diameter in cm, *p*_*S*_ is the calibrated parameter for the tree size mortality (scaling coefficient), and *r*_*S*_ is the exponent that determines the rate at which mortality increases with *d* in the canopy.

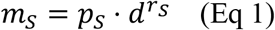

A growth rate-dependent mortality for the upper canopy layer was formulated as a function of biomass increment to account for a higher mortality rate of fast-growing trees, using a logistic function (Eq 2). *ΔB* represents biomass increment in kg C, *p*_*GR*_ is the calibrated parameter for the growth rate mortality, *a* is a correction coefficient for the function, and *r*_*GR*_ is the rate at which mortality increases with *ΔB*.

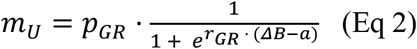

To evaluate the influence of the shape of the mortality formulations (Eq. 2 and 3), we set three mortality rates for each mortality formulation (*r*_*S*_ = 1.5, 2.5 *or* 4 and *r*_*GR*_ = −0.5, −0.8 *or* − 1.4), where the different parameter values control the shape of the curve for low, medium and high curvature. The mortality parameters were calibrated using a cost function described in section 2.2.3.

The same understory mortality was applied to both model setups, with higher mortality rates for the smaller and younger understory cohorts (Eq 3). This equation was adapted from Weng et al. 2015, where *d* is the diameter in cm, *p*_*U*_ is the calibrated parameter and *a, b* are correction coefficients for the understory mortality.

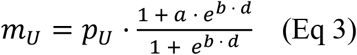

#### 2.2.3. Model calibration

A calibration at the ecosystem-level aggregate was done such that the model was able to realistically simulate average ecosystem photosynthesis and biomass and adequately describe forest dynamics, largely representative of the Swiss forest data used here. We calibrated the model using data from the Lägeren site (CH-Lae), a mixed mountain forest located on the Swiss Plateau at 800 m asl. Data for this site was obtained from the Long-term Forest Ecosystem Research (LWF) project (Thimonier *et al*. 2010). A direct calibration was first done from observations and included leaf mass per unit area (LMA, kg m^-2^), wood density (kg m^−3^) and species-specific allometry parameters (Forrester *et al*. 2017). Calibration was then performed for five model parameters determining the root-shoot ratio, the maximum leaf area index (LAI) limited by light, a scalar for plant respiration and the mortality parameters for each formulation (*p*_*U*_, *p*_*S*_ and *p*_*GR*_). The calibrated targets included mean annual ecosystem-level Gross Primary Productivity (GPP) obtained from the FLUXNET 2015 dataset for Lägeren (Pastorello *et al*. 2020) and the 95^th^ percentile of the LAI in the peak growing season (from June to August) obtained from MODIS (Myneni *et al*. 2015). We used the LWF data to estimate stand-level biomass (kg ha^-1^) and tree size distributions binning five size classes with an approximately equal number of trees per ha. Parameters were calibrated by minimising the root mean square error (RMSE) between the observed and modelled targets. The cost function was defined with equal weighting of errors in all calibrated targets (GPP, LAI, stand biomass and 5-class tree size distribution). Calibration was performed using the Generalised Simulated Annealing algorithm from the *GenSA* R package (Xiang *et al*. 2013).

#### 2.2.4. Simulations

All simulations were initiated with 0.05 saplings per m^2^ and a single plant functional type (PFT) representing a dominant temperate deciduous tree, such as *Fagus sylvatica*. Simulations were run for 1500 years in total, with a spin-up of 700 years to reach steady-state pool sizes. We used temporally constant model forcing data based on meteorological and CO_2_ information obtained from CRU TS (Harris *et al*. 2020) and FLUXNET2015 (Pastorello *et al*. 2020) via the *ingest* R package (Stocker 2020a). Forcing variables include air and soil temperature (°C), precipitation (mm), radiation (W m^−2^), atmospheric pressure (kPa), CO_2_ (µmol mol^−1^), wind speed (m s^−1^), relative humidity (%), and soil water content (%). The LM3-PPA implementation in the *rsofun* R package (Stocker 2020b) was used.

To simulate growth enhancement, the photosynthetic light use efficiency (eLUE) was increased by two levels (+15% and +30%) after the model spin-up. Higher LUE and a resulting tree-level growth enhancement mimics the relief of limitations to carbon assimilation in a generic sense - be it via a growing season extension, enhanced nutrient inputs, relieving reductions of photosynthesis by low temperature, or increasing atmospheric CO_2_. For each mortality model structure (size and growth rate-dependent) and for each curve shape (*r*_1−3_), we ran the simulations for control and the two levels of eLUE.

We evaluated changes in annual total ecosystem-level biomass production (*B*, kg C m^2^), growth (*G*) and mortality (*M*) over time. Note that here, *G* and *M* are defined as fluxes of C (kg C m^2^ yr^-1^), thus differing from tree-level growth, commonly expressed as an increment of diameter per unit of time, or mortality defined as a fraction of dying trees per unit of time. We then calculated the relative changes in total biomass (*dB*/*B*), mortality (*dM*/*M*), carbon turnover rate (*dk*/*k*), and longevity (*dL*/*L*) by comparing *G, M*, and *B* averaged over 600 years before and after the step increase in LUE, and evaluated their ratio with respect to relative changes in growth (*dG*/*G*). The carbon turnover rate *k* (yr^-1^) was defined as the ratio between *M* and *B*, i.e., the inverse of carbon residence time *τ* (yr; *k* = 1/*τ*). Longevity was defined here as the age of the oldest cohort present within the tile.

We also quantified the changes in the self-thinning trajectories resulting from eLUE conditions. We selected the last 600 years of the simulations to ensure that the steady state had been reached. We tested if the STLs were influenced by the levels of LUE by fitting linear models (LMs) with stand density (log-scale) as the dependent variable, and QMD (log-scale) and LUE (as a proxy of growth enhancement) as predictors. The residuals of the models were checked for normality and homoscedasticity.

#### 2.2.5. Evaluation

To evaluate the performance of the model and the overall simulation results, we first ran a sensitivity analysis for the allometric scaling parameter relating diameter and biomass in order to test the changes in the G-B relationship. We used a species-specific allometry parameter for European beech (*θ*_*BM*_ = 2.36 in the LM3-PPA model) to run the simulations and considered a plausible range of parameter values (*θ*_*BM*_ = 2.20, 2.36, 2.50) to test the sensitivity of the G-B relative changes. Second, we evaluated the simulated relationship between growth rate and mean age of canopy trees with respect to the expected negative growth rate-longevity relationship found in the literature (Bigler and Veblen, 2009; Manusch et al., 2012). Cohort-level simulations for each mortality assumption (structure and curve shape) and each LUE level were used to estimate mean growth rate and age of the canopy trees. We selected the two tallest cohorts to estimate the mean age of the canopy trees as a proxy of life expectancy. Mean growth rate was calculated for those cohorts as the diameter increment (mm yr^-1^) averaging across transient years.

## 3. Results

### 3.1. Observational changes in the self-thinning relationships

The STL shifted upward over time (Fig. 1A), i.e., for a given QMD, stands tend to have become denser through time. This emerges from the patterns over time and across sites, thus indicating that the relationship between biomass and density has not been stationary but has shifted significantly (*P*<0.001) over the past decades (see Table S2). The upward shift of the STL over time, i.e., the average increase in density for a given QMD, was ≈0.03-0.04% per *year* for the 55^th^, 75^th^, and 90^th^ percentiles. Unmanaged Swiss forests also exhibited a change in the STL when trees grow more vigorously (Fig. 1B). The LMMs identified a significant (*P*<0.001) positive effect of *growth anomaly* on the intercept of the STL (Table S2). The average increase in density for a given QMD was ≈0.80-1.17% per unit of *growth anomaly* (kg m^2^ yr^-1^) for the 55^th^, 75^th^, and 90^th^ percentiles. The inclusion of both predictors improved model performance based on lower AIC (−539.50<-512.23 for *year* and -305.86<-299.63 for *growth anomalies*). The percentage of stand density variance explained by the fixed effects, i.e., the marginal pseudo-R^2^, ranged from 86 to 87% for both models.

**Figure 1.**
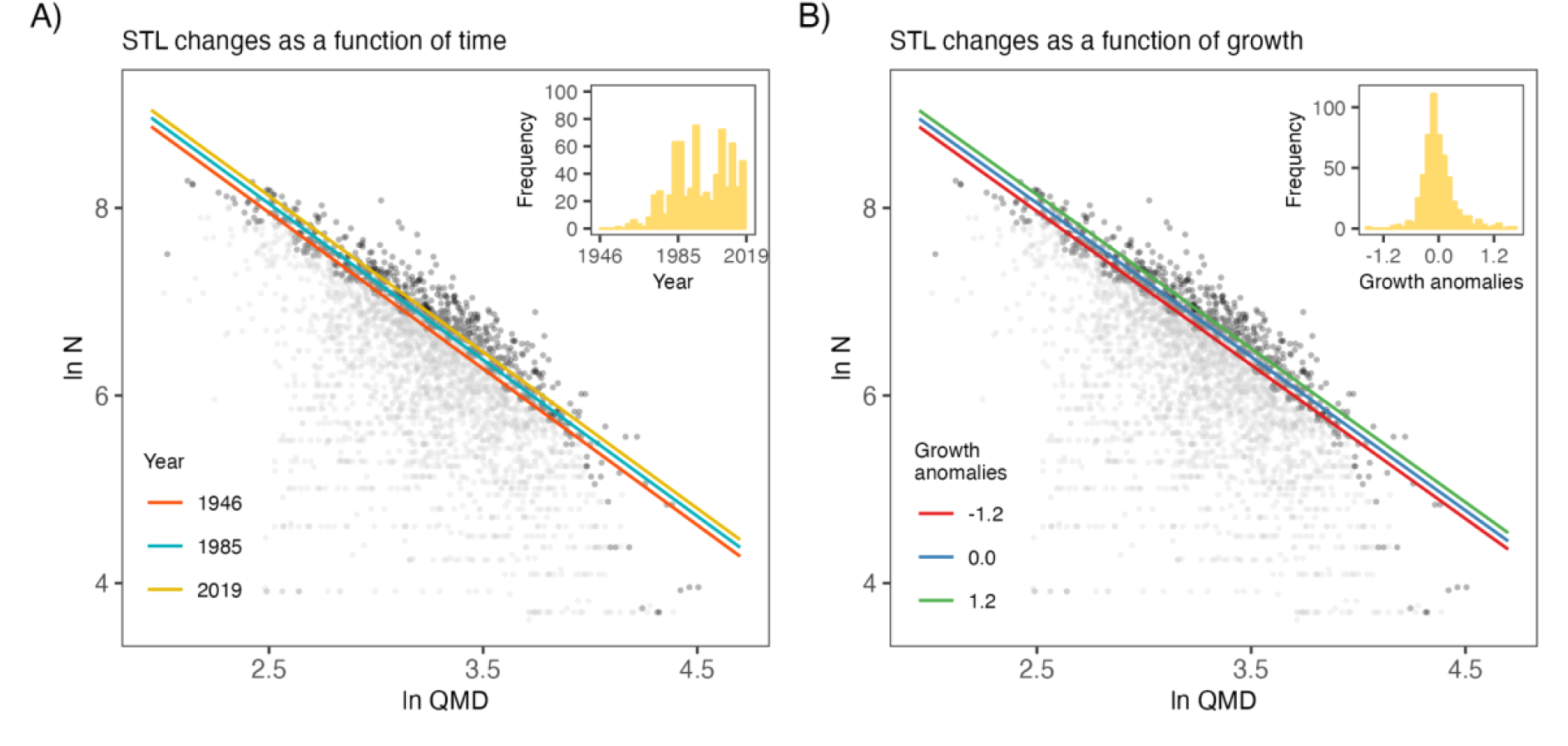
Stand density (N, trees ha^-1^) (log-scale) as a function of QMD (cm) (log-scale) and (A) calendar year, or (B) growth anomalies over the study period for the plots of the pooled NFI, EFM and NFR networks. Dark grey points represent data from plots selected within the 75^th^ percentile and used for model fitting; light grey points are the observations below the selected criterion. Coloured lines represent the fitted STLs. The embedded panels display the distributions of (A) calendar year and (B) growth anomalies for all forest data.

### 3.2. Simulated changes in biomass due to growth enhancement

In response to a 15% (30%) increase in LUE, *G* increased by 17% (35%) on average across the last 600 simulation years before and after the step increase. The higher stimulation of G compared to LUE is due to allocation to woody biomass in the model. *B* increased in response to *G* enhancements in all model setups, irrespective of the mortality structure and shape of the mortality parameterization (Figs. 2A-D and 3A-D). However, the magnitude of *B* varied systematically with the structure and shape of the mortality formulations. Following the size-dependent mortality parameterization, *B* increased by 9-14% (19-28%), and the higher the curvature of the mortality parameterization, the lower the increase in *B*. Following the growth-rate dependent mortality parameterization, *B* increased by 17% (33%), whereas the curvature of the mortality parameterization had no effect on *B*. Once a new steady state of biomass stocks had been attained in the simulations, *M*, expressed in units of living biomass loss per unit area and time, attained the same average level as *G* in all simulations and model formulations. This is a direct consequence of mass conservation but also indicates that under environmental changes and gradually increasing *G, M* increases in parallel, albeit with a lag. The considerable temporal variations of *B* and *M* reflect forest stand dynamics under dynamic equilibria before and after the step increase in eLUE and, consequently, growth.

**Figure 2.**
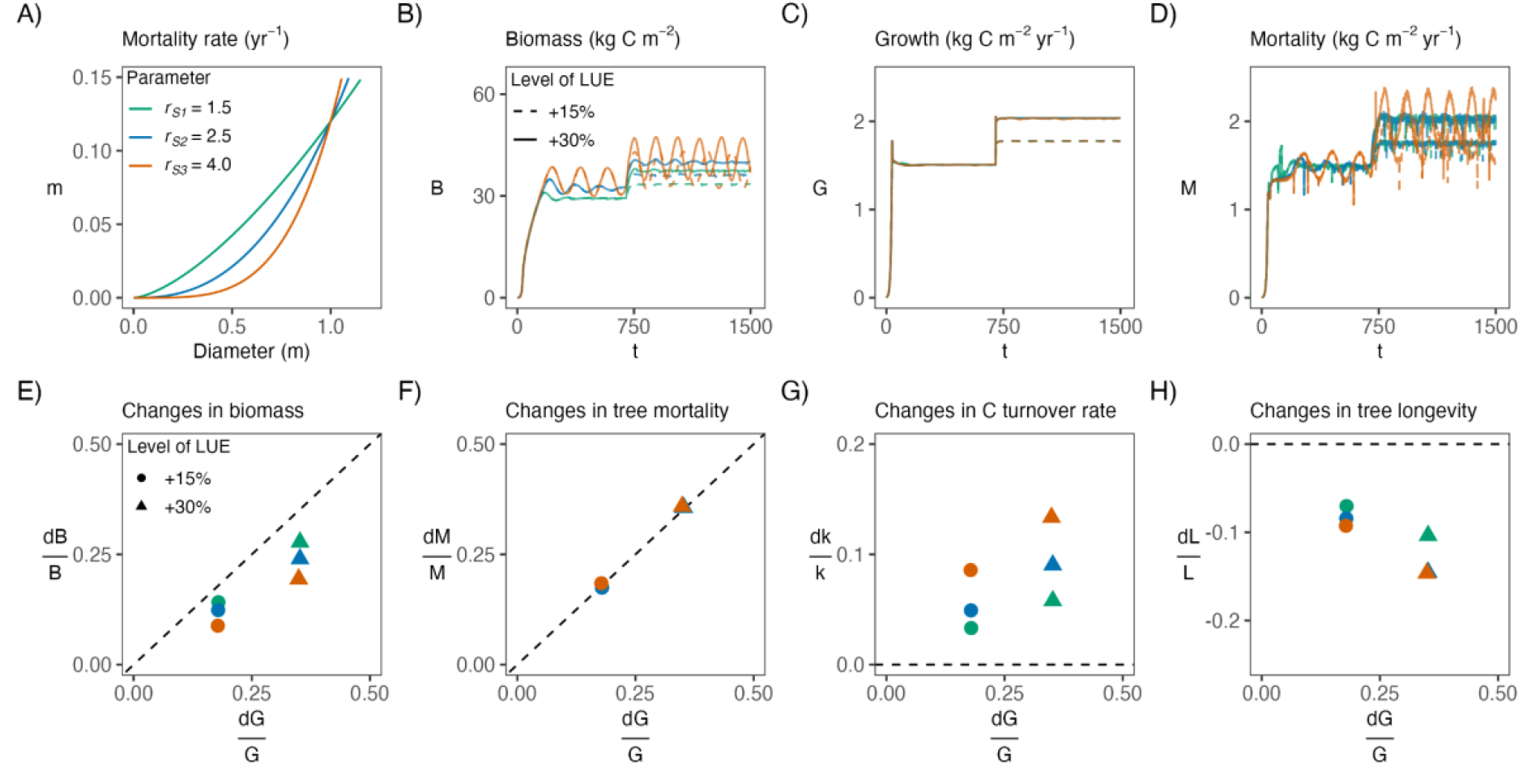
Model simulations for size-dependent mortality and different mortality shapes (A) showing the absolute changes in biomass (B), growth (C) and mortality (D) over time and the relative changes in biomass (E), mortality (F), carbon turnover rate (G) and longevity (H) with respect to relative changes in growth. Colours show the mortality shape (low to high curvature), and line types/point shapes show simulated increases in LUE (15% and 30%). Dashed lines following the 1:1 line (E, F) or the zero-value (G, H) represent the hypothetical *constant turnover rate*.

**Figure 3.**
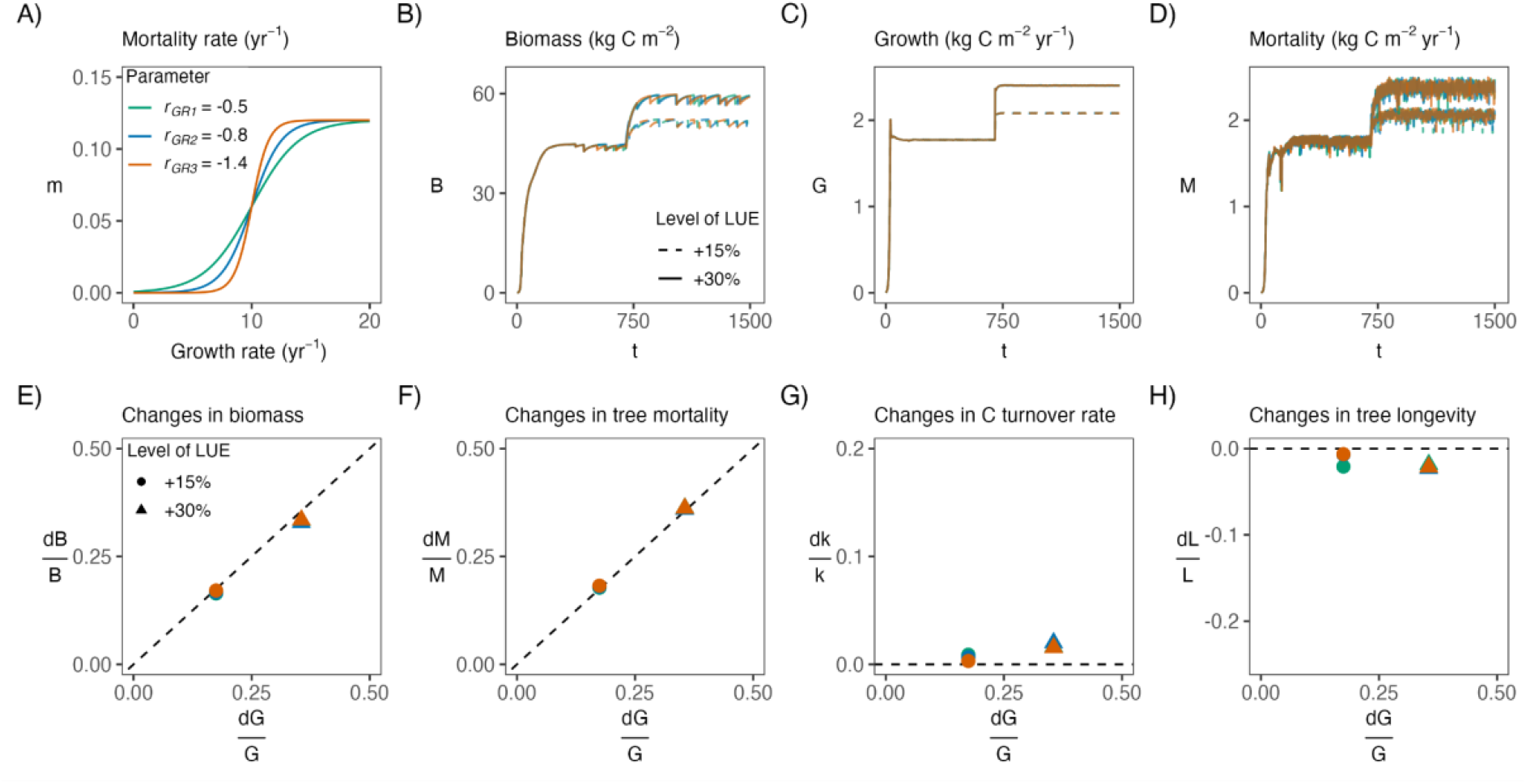
Model simulations for growth rate-dependent mortality and different mortality shapes (A) showing the absolute changes in biomass (B), growth (C) and mortality (D) over time and the relative changes in biomass (E), mortality (F), carbon turnover rate (G) and longevity (H) with respect to relative changes in growth. Colours show the mortality shape (low to high curvature), and line types/point shapes show simulated increases in LUE (15% and 30%). Dashed lines following the 1:1 line (E, F) or the zero-value (G, H) represent the hypothetical *constant turnover rate*.

Comparing the relative increases of different variables to the relative increase in growth yields insights into the (non-) linearity of the system representing forest dynamics and biomass stocks (Box 1). Although *B* generally increases in response to increases in *G* - irrespective of mortality parameterizations - the relative increase in *B* is always smaller than the relative increase in *G* for the size-dependent mortality formulation. The ratio of the respective relative changes varies substantially depending on the curvature of the mortality parameterization (Fig. 2E). This indicates a distinct non-linearity in the system, introduced by the link between *G* and B, and illustrates the degree of this non-linearity (deviation from the 1:1 line in Fig. 2E) is governed by the curvature (non-linearity) of the mortality parameterization as a function of tree size. The growth-rate dependent mortality formulation does not introduce such non-linear behaviour, and the relative increase in *B* is almost identical to the relative increase in *G* (Fig. 3E). Reflecting mass balance constraints, relative increases in *G* and *M* are always identical, irrespective of the structure and shape of the mortality parameterization (Fig. 2F and 3F).

Substantial variations in the relative increases in *B* for a given relative increase in *G* are reflected by the relative changes in the turnover rates and maximum tree longevity. Using the size-dependent mortality formulation, turnover rates increased (Fig. 2G) and maximum tree longevity decreased (Fig. 2H), irrespective of the curvature of the mortality formulation. Using a low curvature parameter of the mortality function (*r*_s1_ = 1.5 in Fig. 2A), smaller relative changes in carbon turnover rates and tree longevity were simulated in response to growth enhancements than when using a pronounced curvature. The highest curvature showed the highest relative increase in turnover rates and the strongest relative decrease in tree longevity in response to growth enhancements. No changes in turnover rates and only small changes in tree longevity were found in response to growth enhancements when mortality was a function of growth rate, independently of the shape of the mortality parameterization (Figs. 3G and 3H). Overall, relative increases in turnover rates were smaller than relative increases in *G*, thus leading to a positive response of *B* in all model setups.

Taken together, the model simulates an acceleration of forest dynamics and a shortening of tree longevity when using a size-dependent mortality formulation. This is measured by the increase in turnover rates, which can be seen along with a reduction of the carbon residence time due to the speeding up of the life cycle (Fig. S2A). As trees grew faster, tree size distributions shifted towards larger sizes (Fig. S3A), despite the reduction in their longevity. This acceleration of forest dynamics did not preclude an increase in steady-state biomass stocks - irrespective of the assumptions regarding the mortality parameterization. If tree mortality was assumed to increase as a direct consequence of faster growth, as embodied in the growth rate-dependent mortality formulation, only slight decreases in carbon residence time were found (Fig. S2B) and trees reached larger sizes before death (Fig. 3B). These simulations did not yield a reduction of tree longevity, and growth enhancements translated directly and nearly linearly into biomass enhancements.

### 3.3. Simulated changes in the self-thinning relationships

Regardless of the mortality formulation, eLUE simulations led to an upward shift in the STL (Fig. 4), suggesting a significant change in the maximum stand density and pointing to larger trees for a given stand density or denser stands for a given average tree size. Biomass was largely constant along the STLs and thus, an upward shift of the STL indicated that biomass increased at conditions where self-thinning is acting. Further, our results revealed the influence of the mortality structure and parameterization on the degree to which the STL is shifted in response to growth enhancements. Size-dependent mortality with a flatter shape predicted a stronger increment of stand density for a given QMD (≈2% for 15% eLUE and ≈3% for 30% eLUE, Fig. 4A), while functions with a higher curvature led to a weaker change in the STL (≈1% for 15% eLUE and ≈2% for 30% eLUE, Figs. 4B and 4C). For the growth rate-dependent mortality formulation, the STL had a significant increase in the intercept, indicating that stands support higher densities when increasing growth, independently of the shape of the curve (≈2% for 15% eLUE and ≈3% for 30% eLUE, Figs. 4D-F).

**Figure 4.**
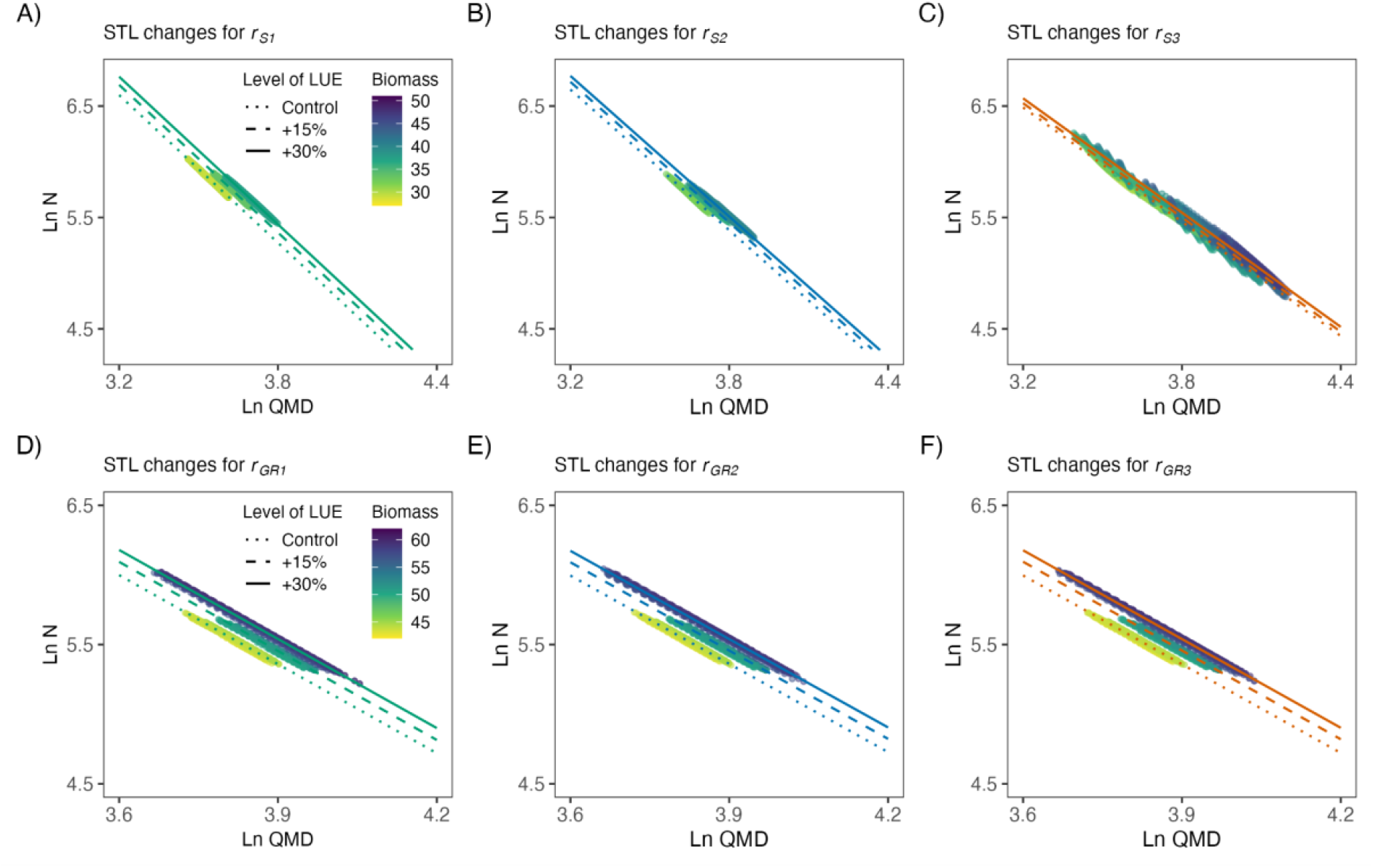
Simulated relationships between stand density (N, trees ha^-1^) and quadratic mean diameter (QMD, m) for each alternative mortality formulation: (A-C) size-dependent, (D-F) growth rate-dependent and each curvature shape (*r*_*1*_-*r*_*3*_, line colours) under simulated increases in LUE (line types). The log-log slopes of the regression lines represent the maximum stand density. Points are the simulated data for each combination coloured as a function of total biomass (kg C m^−2^).

### 3.4. Evaluation of model performance

The sensitivity analysis of the allometric scaling parameter that related diameter and biomass confirmed the positive relationship between growth and biomass, for all the plausible values tested (Fig. S4). For all values of *θ*_*BM*_ (2.20, 2.36 and 2.50), the size-dependent mortality led to a non-linear *G-B* relationship (Figs. S4A-C). The growth-rate dependent mortality had a linear behaviour for all the parameter values considered, with relative increases in *G* being similar to increases in *B* (Figs. S4D-F). In both mortality structures, the higher the *θ*_*BM*_, the stronger the relative increase in *B*.

The relationship between simulated mean growth rates and age of the canopy trees for the different mortality assumptions is shown in Fig. S5. The size-dependent mortality featured a strongly negative relationship between growth and life expectancy (Fig. S5A), independent of the shape of the mortality curve (*r*_*p*_ = [-0.99, -0.95]). For these simulations, enhanced growth rates lead to shorter life expectancy. The growth rate-dependent mortality also featured a negative correlation between mean growth and age of canopy trees (Fig. S5B), which varied depending on the curve shape (*r*_*p*_ = [-0.97, -0.80]). In this case, there are at least two opposite factors affecting the mean age of the canopy trees. If trees grow faster, they will have a high mortality rate. However, the large-sized trees prevent the young trees from reaching the top layer. If trees grow slower, they will have a low mortality rate and thus higher longevity, but younger trees get into the upper canopy. The shape of the mortality curve may play a role in determining the equilibrium state, i.e., the mean age at equilibrium tree compositions.

## 4. Discussion

We combined forest observations and model simulations to evaluate to what extent tree growth enhancements lead to persistent increases in forest biomass stocks. We found that the position of the STL has shifted upwards over time and as growth rate increased in unmanaged, closed-canopy forests in Switzerland. A biomass increase under enhanced growth was simulated in all model setups, but the magnitude of the change varied substantially depending on the shape of the mortality parameterization. The relative changes in biomass were smaller than the relative changes in growth, indicating a reduction in the apparent carbon residence time and in tree longevity. The increase in steady-state biomass with enhanced growth was also reflected in the upward shift of the modelled STL - consistent with observations.

### 4.1. Growth enhancements lead to biomass increments

Irrespective of the mortality assumption, a positive net increment in biomass was modelled despite the reduction in carbon residence time and tree longevity. When mortality was size-dependent, our study framework indicated a non-linear *G-B* relationship, as described by the *accelerated turnover* response (Box 1). These findings suggest a trend towards higher densities per unit of tree size and reveal that increasing biomass stocks and decreasing C residence times are not mutually exclusive. This reconciles reports of tree longevity reductions (Bugmann & Bigler 2011; Büntgen *et al*. 2019; Brienen *et al*. 2020) with model predictions of increased forest biomass (Terrer *et al*. 2019; Yu *et al*. 2019; Pugh *et al*. 2020), both of which are consistent with the mechanistic understanding developed here. When mortality was defined in terms of growth rate, our results showed an almost linear *G-B* relationship, reflecting a *constant turnover rate* response (Box 1). This yields only a small decrease in longevity and carbon residence time, as has been represented in models that account for a constant background mortality (Bugmann *et al*. 2019). Notably, none of the mortality assumptions implemented in the model, nor the data suggested a *constant self-thinning* response (Box 1), as underlined in the GFDY hypothesis. Yet, the ratio of relative changes in growth and biomass was critically affected by the shape of the mortality formulations. As we show here, the stronger the curvature in the size-mortality parameterization, the smaller the increase in biomass and the smaller the upward shift in the STL. There is still uncertainty about model structural choices, and different assumptions and parameterizations may lead to contrasting results. For example, contrasting results by Brienen *et al*. 2020 indicated a lack of long-term biomass increments in response to a temporal trend towards increased growth. This is possibly related to their choice of a highly non-linear size-mortality parameterisation, fitted to data that reflects a growth-longevity relationship across species - not a temporal relationship that underlies the forest inventory data analysed here.

Carbon assimilation rates in terrestrial ecosystems have increased steadily as atmospheric CO_2_ concentrations have risen over the past century (Campbell *et al*. 2017; Walker *et al*. 2021). In parallel, rising temperatures have led to an expansion of the growing season in winter-cold climates (Piao *et al*. 2019). Simultaneously, a substantial terrestrial C sink has persisted (Keeling *et al*. 1996; Friedlingstein *et al*. 2021). Yet, gains in carbon storage, driven by increased photosynthesis and growth, have been argued to be transitory (Körner 2017), and ultimately limited by other resources (e.g., nutrients) and negative feedbacks arising through forest dynamics. These mechanisms linking changes in terrestrial photosynthesis and C storage remain uncertain (Huntzinger *et al*. 2017) because a multitude of processes and feedbacks are involved at different scales, ranging from leaves to trees, forest stands, ecosystems, the landscape, and the globe (Walker *et al*. 2021; Maschler *et al*. 2022). The *G-B* relationships and the STL shifts described here are relevant for the C cycle dynamics and the propagation of effects by increased photosynthesis and growth to the scale of a forest stand. How the processes are represented in vegetation models will determine the accuracy of predictions of forest responses under elevated CO_2_ and other environmental changes (Andresen *et al*. 2016; Davies-Barnard *et al*. 2020; Bugmann & Seidl 2022). With the advent of demographic representations in global vegetation and terrestrial carbon cycle models, there is a need for constraining alternative process representations with observations. Novel cohort-based vegetation demography models (Fisher *et al*. 2018), such as the LM3-PPA, resolve tree age and height structure and enable a more mechanistic treatment of forest dynamics and tree mortality. This yields a mechanistic foundation to project responses to environmental change and enables globally distributed forest inventory data to be used for constraining the models. However, observations are sparse due to the long timescale of forest demographic processes. The approach taken in this study enabled us to test the GFDY hypothesis via the STL changes observable from data that inform the unobservable (simulated) steady-state biomass response to growth enhancement.

### 4.2. Non-linearity in the growth-biomass relationship

The ratio of relative changes of growth and biomass yields insights for characterising carbon cycle dynamics in forest models and normalises effects with respect to absolute magnitudes of simulated biomass and growth. Our study shows that this ratio is subject to the representation of mortality in the model. In LM3-PPA, the PPA warrants that the tree crowns fill gaps in the canopy through phototropism (Purves *et al*. 2008). In our simulations, a growth enhancement skews the distribution of trees to larger sizes, decreases the number of trees in the canopy, and increases tree numbers in the understory. Under conditions of higher growth, this replacement is accelerated, leading to higher mortality rates, lower longevity, and a subsequent decrease in the carbon residence time (Needham *et al*. 2020). We tested the sensitivity to mortality parameters and model structural choices. However, other processes affecting resource accessibility to tree individuals and their neighbours may influence the non-linearity of the *G-B* relationship. This includes parameters regarding allometric scaling, height-dependent crown organisation and light penetration in the canopy. We additionally evaluated the influence of alternative allometric scaling parameters. This indicated that the finding of generally positive biomass changes in response to growth increases is robust against a wider choice of model formulations and parameterizations (see Fig. S3).

Understanding the causes of observed mortality trends will help to improve the way mortality is included in vegetation demography models, which is critical for accurate projections of global terrestrial carbon storage (Friend *et al*. 2014). Different modes of mortality (e.g., hydraulic failure, carbon starvation) could be incorporated into models and may lead to different ratios of relative changes. Different model structural assumptions regarding light distribution cannot be easily tested within a single modelling framework. Future work including model intercomparisons to test simulations with a set of alternative models would be helpful in informing the generality of the positive *G-B* relationship found here. Importantly, to evaluate model reliability in accurately simulating *G-B* links, a focus has to be set on whether they capture self-thinning relationships (slope, position, and their change over time) as suggested by the data. Thus, combined analyses of models and forest observations will be needed to project how changes in environmental conditions will affect competition for resources and forest dynamics in a future climate (McDowell *et al*. 2018). A large number of long-term monitoring forest demographic rates are required to better understand the links between growth and biomass and to constrain influential, yet not directly observable model parameters.

### 4.3. Endogenous and exogenous factors affecting carbon residence times

It is of crucial importance to distinguish between changes in carbon residence times caused by endogenous (i.e., growth, density-driven mortality) and exogenous factors (e.g., climate, climate-driven disturbances). Here, we focused on the former. Observations from tropical forests have suggested that increases in productivity combined with persistently higher mortality led to shorter carbon residence times (Brienen *et al*. 2015; Hubau *et al*. 2020). Still, no clear consensus exists about the trade-offs between growth and tree longevity and their temporal changes within species (Cailleret *et al*. 2017). Determining the growth-lifespan trade-offs under current environmental conditions is subject to constant growth conditions and resource availability. However, environmental changes affect growth conditions for all species and may relieve constraints shifting the trade-offs, as suggested by our results. Changes in tree mortality are also caused by changes in the environment (DeSoto *et al*. 2020). Disturbances are becoming more frequent (Sommerfeld *et al*. 2018), leading to enhanced tree mortality around the world (Senf *et al*. 2018).

Evidence suggests that carbon residence times in forest biomass have reduced in the past (Yu *et al*. 2019) and may be reduced by future climate change. Rising temperature, vapour pressure deficit (VPD) levels and more frequent drought episodes (Schwalm *et al*. 2017; McDowell *et al*. 2020) can reduce photosynthetic C uptake as trees close their stomata to prevent hydraulic failure. This may cancel any potential benefit from elevated atmospheric CO_2_, leading to lower growth (Yuan *et al*. 2019) and higher mortality (Park Williams *et al*. 2013). Climate-driven risks may thus lead to higher competition for water and override growth-related forest density trends (Anderegg *et al*. 2020). Our findings highlight that growth enhancement causes simultaneous increases in biomass and decreases in carbon residence times and tree lifespans and the non-linearity in the growth-biomass relationship is to be understood as representing effects within species.

### 4.4. Interpreting self-thinning relationships

The non-linear modelled growth-biomass relationship is consistent with the empirical results suggesting temporal trends in the STL and a link to growth variations across plots and time. We applied the STL concept to mixed, often uneven-aged forests in Switzerland to detect whether constraints governed by density-driven mortality have been relieved. Traditionally, the focus of the STL has been restricted to even-aged monospecific stands, and the power-law exponent (i.e., the slope of the STL) was proposed to be constant and universal (Reineke 1933; Yoda 1963; Westoby 1984). Further studies showed that the STL directly reflects allometric and metabolic scaling, linking tree size, stand structure and biomass stocks (Enquist *et al*. 2009). Generally, higher intercepts and slopes are associated with fertile soils (Morris & Charles Morris 2003; Bi 2004), which are able to reach higher densities (Weiskittel *et al*. 2008; Charru *et al*. 2012).

Self-thinning dynamics have also been described in mixed forests (Midgley 2001; Mrad *et al*. 2020) and the application has been generalised to multispecific stands (Rivoire & Le Moguedec 2012; Forrester *et al*. 2021). The self-thinning relationship emerges from density-driven mortality due to resource competition between individuals, neglecting mortality due to external factors. Our approach excluded areas under management, and we selected plots from the upper quantiles (featuring high density for a given QMD) as those subject to self-thinning. By doing so, we ensured to remove or at least minimise external effects from natural or anthropogenic past disturbances. Indeed, we found a clear negative linear relationship as seen in pure even-aged stands and we determined the stands where the STL has been reached with the upper edge of the point cloud.

The STL approach allowed us to control for stand age effects on biomass, thus revealing shifts in biomass storage without having to rely on observations of mature stands. Our empirical analyses suggested a tendency toward denser stands for a given QMD over time and indicate that stand density is related to growth vigour. These results are consistent with empirical evidence from Kubiske *et al*. 2019 who reported increasing intercepts of the STLs under higher CO_2_, with the consequent higher stand biomass levels in the long term. Recent findings also indicate that climatic variables (Brunet-Navarro *et al*. 2016; Forrester *et al*. 2021) influence the STL, although other studies found that it remained constant over time (Pretzsch *et al*. 2014). Importantly, the STL in mixed forests can also change when the relative proportion of species changes (Reyes-Hernandez *et al*. 2013), e.g., due to succession. However, the Swiss forest stands used in our analyses did not feature strong changes in species composition according to the Bray-Curtis dissimilarity index (see Table S1). Our analysis also considered species effects by including the dominant species per plot as a random factor to control for species composition.

Further, our framework of evaluating changes in the STL in observations and simulations (with one PFT) avoids confounding effects to the largest extent possible. Our findings confirm that STLs are not static, simply reflecting edaphic factors, but are changing with changes in the environment. This is relevant for forest management, which often relies on the STL to inform wood harvesting and plantation management (Nagel *et al*. 2017). Importantly, the slope of the STL may also change as forest stands mature. Assuming a stationary self-thinning trajectory and a steeper slope as stands mature (constant final yield), would imply a downward shift of the fitted STL. Future work needs to investigate if shifts in these relationships also occur in primary forests along broader environmental gradients. Confronting modelled with empirical relationships will enable new insights into the links between forest dynamics and biomass.

## 5. Conclusions

Forest responses to global environmental changes are still unclear and difficult to study due to multiple interactions and anthropogenic disturbances. We focused on the mechanisms of forest stand dynamics and demography that determine the link between changes in tree growth and stand-level biomass stocks. We find that unmanaged closed-canopy forests in Switzerland have become denser for a given size over the past six decades, and we identify a positive relationship between growth and stand density. These observations are consistent with simulations showing that growth enhancement leads to increases in forest biomass and changes in the self-thinning relationship. However, relative changes in biomass are smaller than relative changes in growth, indicating an apparent reduction in carbon residence time. We show that this effect critically depends on the shape of the mortality parameterization. This data-supported mortality modelling yields new insights into the causes of currently observed terrestrial carbon sinks and future responses. Our study provides a better understanding of whether and how growth enhancements drive higher C storage - a key open question in carbon cycle research and highly relevant in the context of climate and Earth system changes.

## Acknowledgements

We gratefully acknowledge the data providers and their long-term work to maintain and measure the different forest plots network. LM and BDS were funded by the Swiss National Science Foundation grant no. PCEFP2_181115. We acknowledge WSL and ETH and their scientists, technicians and data managers who designed, carried out and maintained the measurements on the permanent monitoring plots used in this study. Model calibration was based on data from the Swiss Long-term Forest Ecosystem Research (LWF). Data analyses and evaluations were based on data from (a) the Swiss National Forest Inventory (NFI), (b) the Experimental Forest Management (EFM), and (c) the Natural Forest Reserves (NFR). The Swiss Forest Reserve Research Network is supported by the Swiss Federal Office for the Environment (FOEN), WSL, and ETH Zurich. This work is a contribution to the LEMONTREE (Land Ecosystem Models based On New Theory, obseRvations and ExperimEnts) project, funded through the generosity of Eric and Wendy Schmidt by recommendation of the Schmidt Futures program. BDS acknowledges support from this project.

## Authors’ contributions

BDS and LM conceived the study; BDS, EW and LM developed and implemented the model code; LM calibrated the model, ran the simulations, and conducted the empirical analyses; HB and DIF gave substantial inputs to the study design; DIF provided the EFM dataset; MLH and HB provided the NFR dataset; BR provided the NFI dataset and helped with the variable selection; VT provided the LWF data to calibrate the model. All authors contributed to manuscript development and gave final approval for publication.

## Code availability

Code for the data analysis of this study is available at the GitHub repository DOI: <u>10.5281/zenodo.7326085.</u>

