## Supplementary for "Tree growth enhancement drives a persistent biomass gain in unmanaged temperate forests"

**Content:**

Supplementary Figures S1-S5

Supplementary Tables S1-S2

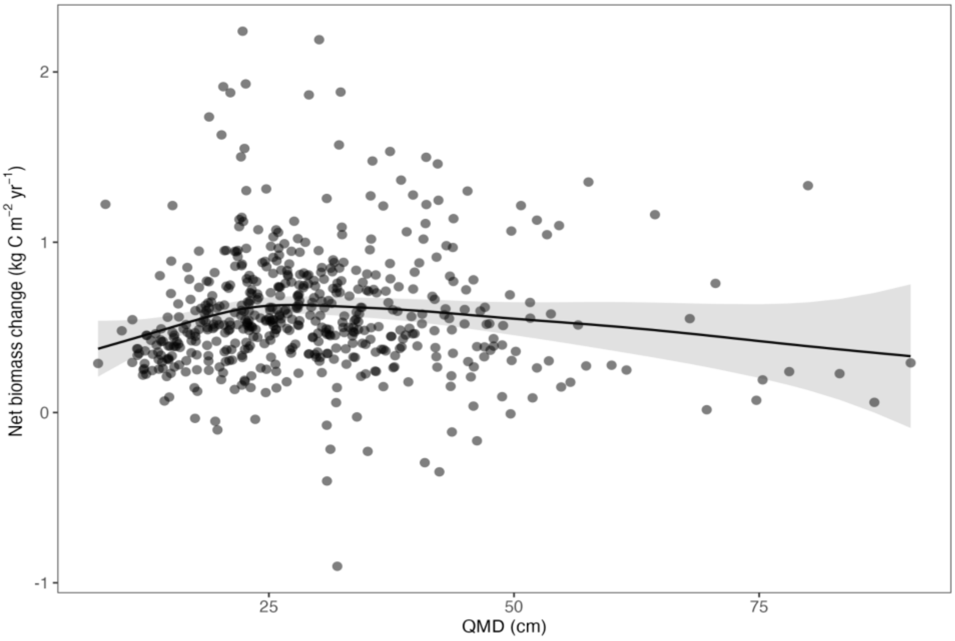


**Fig. S1.** Effect of mean DBH (cm) on net biomass change (kg C $m^{-2}$ $yr^{-1}$) according to the GAMM fitted for the selected stands. The shaded area represents the standard error of the fit. Residuals of the model were calculated as the difference between observed and predicted values.

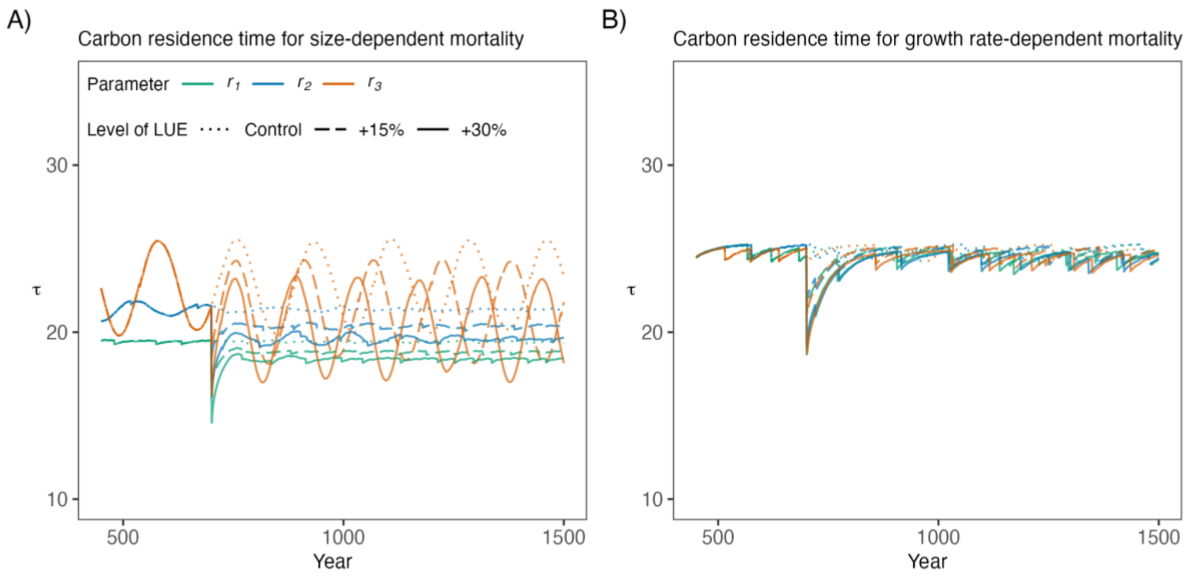


**Fig. S2.** Carbon residence time when mortality is (A) size-dependent and (B) growth rate-dependent. Colours represent the mortality parameterizations (curvature of the function) and line types represent the increases in LUE (control, +15% and +30%).

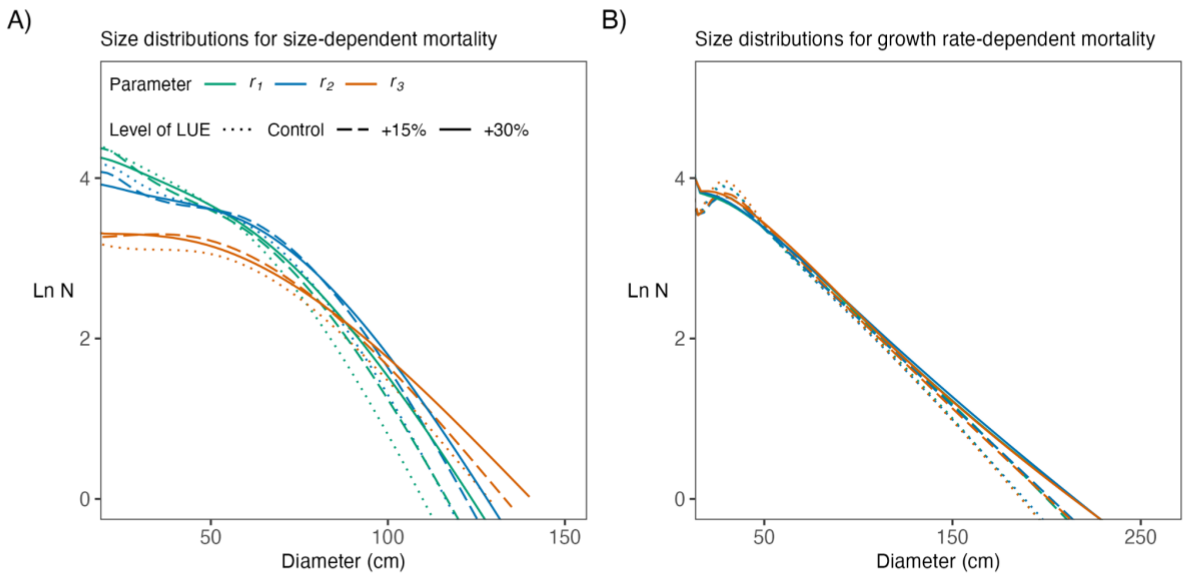


**Fig. S3.** Tree size distributions, i.e., mean number of trees per ha (N, log-scale) and diameter (d, cm) for the last 600 years of simulations in the (A) size-dependent and (B) growth-rate mortality formulations. Colours indicate the shape of the function and line types represent the levels of LUE (control, +15% and +30%).

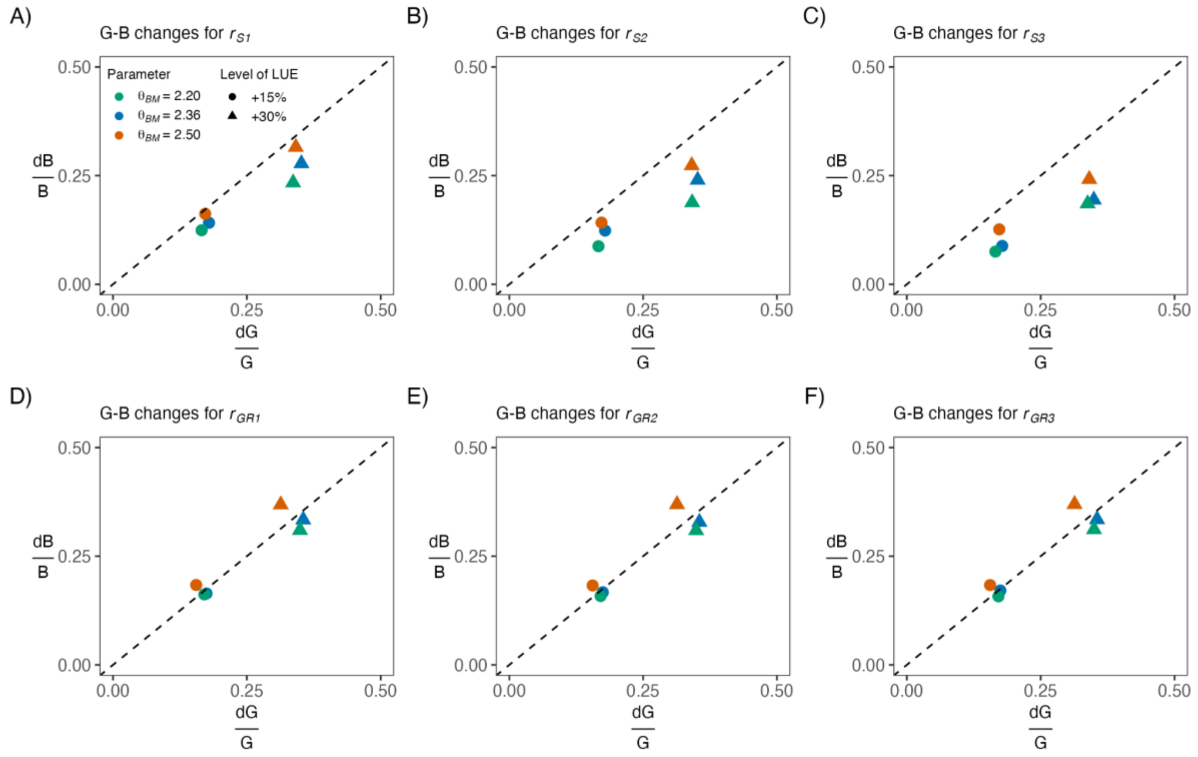


**Fig. S4.** Sensitivity analysis of the scaling allometric parameter relating diameter and biomass in the LM3-PPA model. Panels show the G-B relationships for the (A, B, C) size-dependent and (D, E, F) growth-rate mortality formulations with different structures. Colours indicate the parameter values ($\theta_{BM} = 2.20, 2.36, 2.50$) considered and shapes represent the levels of LUE (+15% and +30%).

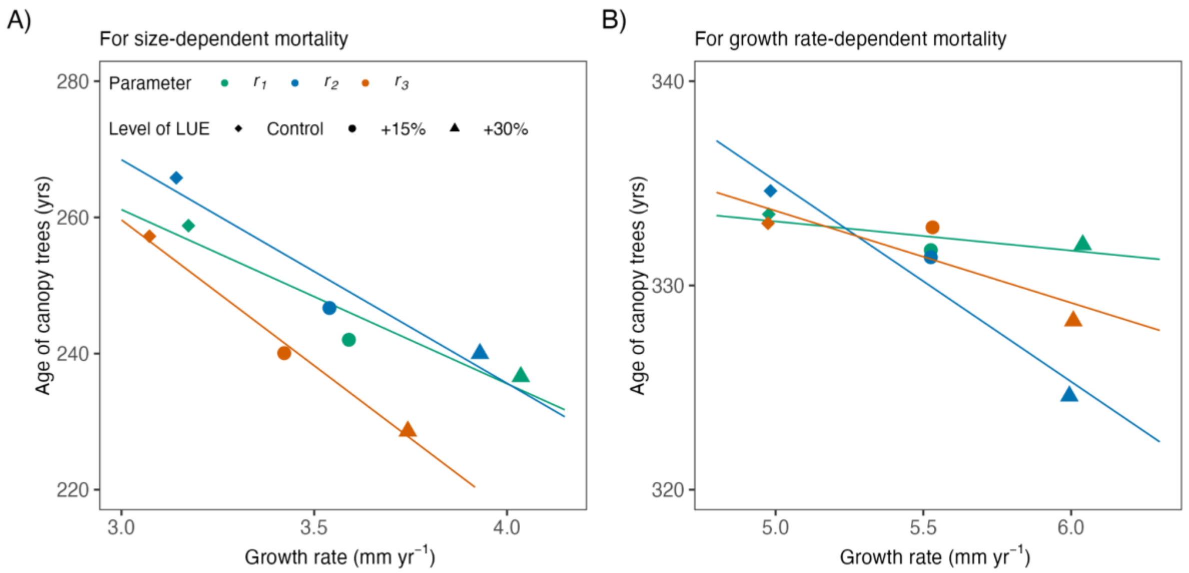


**Fig. S5.** Relationships between mean growth rates and age of canopy trees for the two tallest cohorts after the spin-up years for the (A) size-dependent and (B) growth-rate mortality formulations. Colours indicate the curvature of the function and shape represents the levels of LUE (control, +15% and +30%). Lines are linear regressions for each mortality structure.

**Table S1. Characteristics of the plots for the different monitoring data sources.** Values are means ± SD.

| | NFI | EFM | NFR |
|---|---|---|---|
| Number of plots | 516 | 18 | 269 |
| Plot size (ha) | 0.048 ± 0.006 | 1.15 ± 2.57 | 0.43 ± 0.41 |
| Timespan | 1983-2017 | 1933-2019 | 1955-2019 |
| Measurement intervals (years) | 10 ± 2 | 11 ± 4 | 12 ± 4 |
| Elevation (m a.s.l) | 1430 ± 440 | 1400 ± 538 | 827 ± 428 |
| DBH (cm) | 30.1 ± 11.4 | 27.9 ± 8.9 | 23.0 ± 8.8 |
| Stand density (trees $ha^{-1}$) | 444 ± 329 | 1007 ± 628 | 1203 ± 713 |
| Net biomass change (kg $m^{-2}$ $yr^{-1}$) | 0.37 ± 0.38 | 0.49 ± 0.42 | 0.52 ± 0.25 |
| Bray-Curtis Dissimilarity index | 0.26 ± 0.24 | 0.14 ± 0.11 | 0.20 ± 0.13 |

**Table S2. Main statistics of the LMMs fitted to describe the changes in the self-thinning relationships as a function of calendar year (Year) and growth anomalies ($G_{anom}$).** Variables were standardised, so effect sizes are directly comparable within models. The sensitivity analysis includes results for the 55th, 75th and 90th percentiles of data. $\sigma_u$ denotes the standard deviation of the random intercepts (Plot ID and Dominant species, respectively). The marginal $R^2$ describes the goodness of model fit given fixed effects only, while the conditional $R^2$ describes the goodness of model fit including fixed and random effects. Model parameter significance is annotated as follows: *($p < 0.05$), **($p < 0.01$), ***($p < 0.001$).

| Parameters | 55th percentile | 75th percentile | 90th percentile |
|---|---|---|---|
| Intercept | 6.684 ± 0.024 *** | 6.816 ± 0.031 *** | 6.833 ± 0.055 *** |
| QMD | -0.612 ± 0.011 *** | -0.685 ± 0.013 *** | -0.785 ± 0.023 *** |
| Year | 0.041 ± 0.004 *** | 0.034 ± 0.005 *** | 0.028 ± 0.009 ** |
| $\sigma_u$ | 0.250 / 0.083 | 0.229 / 0.104 | 0.264 / 0.161 |
| Marg. $R^2$ | 0.841 | 0.864 | 0.851 |
| Cond. $R^2$ | 0.980 | 0.985 | 0.992 |
| Intercept | 6.694 ± 0.021 *** | 6.815 ± 0.030 *** | 6.822 ± 0.055 *** |
| QMD | -0.635 ± 0.011 *** | -0.673 ± 0.014 *** | -0.786 ± 0.025 *** |
| $G_{anom}$ | 0.018 ± 0.004 *** | 0.026 ± 0.006 *** | 0.029 ± 0.011 * |
| $\sigma_u$ | 0.239 / 0.065 | 0.222 / 0.095 | 0.239 / 0.157 |
| Marg. $R^2$ | 0.844 | 0.867 | 0.865 |
| Cond. $R^2$ | 0.979 | 0.984 | 0.983 |
